# The mature N termini of *Plasmodium* effector proteins confer specificity of export

**DOI:** 10.1101/2023.05.17.541120

**Authors:** Muhammad M. Hasan, Alexander J. Polino, Sumit Mukherjee, Barbara Vaupel, Daniel E. Goldberg

## Abstract

The intraerythrocytic malaria parasite *Plasmodium falciparum* exports hundreds of proteins into the host red blood cell (RBC). Most are targeted to the ER by a stretch of hydrophobic amino acids and cleaved further downstream at a conserved motif called the Protein Export Element (PEXEL) by the ER protease plasmepsin V (PM V). The mature effectors then travel through the secretory pathway to the parasitophorous vacuole (PV) that surrounds the parasite. There, PEXEL proteins are somehow recognized as export-destined proteins, as opposed to PV-resident proteins, and are selectively translocated out into the RBC. The mature N terminus appears to be important for export. There is conflicting data on whether PM V cleavage is needed for proper export, or whether any means of generating the mature N terminus would suffice. We replaced the PEXEL-containing N-terminal sequence of an exported GFP reporter with a signal peptide sequence and showed that precise cleavage by signal peptidase, generating the proper mature N terminus, yields export competence. Expressing a construct with only the native ER targeting signal without the PM V cleavage site dramatically decreased the amount of a mature PEXEL reporter, indicating that the hydrophobic stretch lacks an efficient cleavage signal. Therefore, the PEXEL motif functions as a specialized signal cleavage site when appropriately located after an ER targeting sequence. Our data suggest that PM V cleavage and RBC export are two independent events for PEXEL proteins. We also tested and rejected the hypothesis that an alpha-helical mature N terminus is necessary for export.

**Importance:** Malaria parasites export hundreds of proteins to the cytoplasm of the host red blood cells for their survival. A five amino acid sequence, called the PEXEL motif, is conserved among many exported proteins and is thought to be a signal for export. However, the motif is cleaved inside the endoplasmic reticulum of the parasite and mature proteins starting from the fourth PEXEL residue travel to the parasite periphery for export. We showed that the PEXEL motif is dispensable for export as long as identical mature proteins can be efficiently produced via alternative means in the ER. We also showed that the exported and non-exported proteins are differentiated at the parasite periphery based on their mature N termini, however, any discernible export signal within that region remained cryptic. Our study resolves a longstanding paradox in PEXEL protein trafficking.

## Introduction

The malaria parasite *Plasmodium falciparum* asexually replicates in the human Red Blood Cell (RBC). An infected RBC is extensively modified by the parasite to make it compatible for growth. Hundreds of parasite proteins are exported into the RBC for this purpose and as such, protein export is critical for parasite survival (1, 2).

Exported proteins are first loaded into the parasite endoplasmic reticulum (ER) and then progress along the secretory pathway by vesicular trafficking to be released outside the parasite plasma membrane (PPM) (3–7). A membrane-bound space called the parasitophorous vacuole (PV) surrounds the PPM throughout the parasite’s replication cycle (8–11). To reach the RBC cytoplasm and beyond, exported proteins, as opposed to PV-resident proteins, must translocate through a multiprotein complex in the PV membrane called Plasmodium translocon of exported proteins (PTEX) (12–16).

In eukaryotic cells, secretory proteins generally have a hydrophobic signal peptide at their N terminus, which targets the protein to the ER membrane. It is then cleaved by the ER protease signal peptidase (SP) to release the secretory protein into the ER lumen. The mature protein can reside in the ER or travel to other organelles depending on the presence of diverse organelle-targeting signals in its sequence. The absence of any such signal generally takes the protein to the plasma membrane to be released outside the cell (17). Similarly, *Plasmodium* proteins with a signal peptide travel to the PV, and for further export into the RBC, an additional signal is necessary (18–20).

A pentameric motif was found to be conserved in most exported proteins of *Plasmodium* and was named the *Plasmodium* Export Element (PEXEL)/Host Targetting motif (21, 22). These proteins are cotranslationally loaded into the ER by a stretch of hydrophobic residues resembling a signal peptide, and the PEXEL motif is generally located after a non-conserved “spacer” region following that signal (5, 23, 24). Minimal constructs capable of export must include around 10 residues downstream of the PEXEL motif as well. However, there is very little primary sequence conservation in these critical residues, and replacing them with a stretch of alanine suffices for export (6, 23, 25).

The PEXEL motif is composed of arginine and leucine in the first and third positions and aspartate, glutamine or glutamate in the fifth position, often expressed as RxLxE/Q/D. Interestingly, the motif is cleaved between the third and the fourth residues by an ER resident aspartic protease called plasmepsin V (PM V), and the mature proteins starting with a comparatively less unique signal of xE/Q/D are then exported (26–28). This raises the question that if the PEXEL motif’s most conserved part (RxL) is cleaved off in the ER, how can it convey export-specificity at the PV? Multiple hypotheses have been offered in this regard. One model is that PM V hands over the cleaved PEXEL proteins to specific chaperones that guide them in the secretory pathway till their delivery to PTEX (27). A recently published study proposed that HSP101, a component of the PTEX translocon, binds to nascent PEXEL proteins at the ER and takes them to the PTEX following PM V cleavage (29). Another model is that PEXEL proteins enter the ER through a PM V-containing translocon that is distinct from the SP-containing translocon, which specifies a selective route to the PTEX complex (4). Mature PEXEL proteins are N-terminally acetylated, which has also been speculated to be a postmark for their eventual export, though acetylation alone is insufficient to achieve export (30, 31). None of these theories has been validated convincingly.

Taking a step back, several experiments have tested the assumption that the PEXEL motif confers export capacity to *Plasmodium* proteins with disparate conclusions. Mutating the conserved R and L, or deleting the PEXEL motif altogether in reporter constructs inhibited their export to the RBC (23, 26, 32). However, these alterations affected the cleavage by PM V, and thus only verified the requirement of the PEXEL motif for cleavage at the ER and not for RBC export per se. More pertinent experiments in this regard would be to check the exportability of mature reporters that are identical to PEXEL-cleaved reporters, albeit in the absence of the PEXEL motif itself. One such construct was designed by removing the region following the hydrophobic stretch to the RxL sequence of a PEXEL protein so that the mature reporter started with the sequence xE after the cleavage by SP at the end of the hydrophobic stretch (27). It did not get exported into the RBC, indicating that the PEXEL motif is required in nascent PEXEL proteins for export. On the other hand, another study tested reporter constructs where the mature parts of PEXEL proteins were preceded by a signal anchor (an ER loading signal, similar to a signal peptide but without the signal cleavage site) and a self-cleaving viral capsid protease sequence that cleaves the reporter right before the xE/Q/D sequences (33).

Interestingly, these reporters were exported into the RBC, suggesting that the signal for export resides downstream of the PEXEL cleavage site and the motif itself is not required for export. Reporters starting with mature N terminal residues of PEXEL proteins were also exported when targeted to the ER by an internal transmembrane domain, supporting the later proposition (34).

In this study, we tested the essentiality of the PEXEL motif for *Plasmodium* protein export with reporter constructs that generate identical mature proteins in the ER, albeit differing in the presence or absence of a PEXEL motif. Our data support a model where the PEXEL motif is required for the cleavage of a protein in the ER but is irrelevant for export to the RBC. Rather, the mature N-terminal domain of PEXEL proteins alone determines their localization once they reach the PV.

## Results

### Serial fractionation can differentiate the localization of PV-resident and RBC-exported protein

We expressed minimal constructs (containing the hydrophobic stretch, spacer, PEXEL motif and short mature N termini of variable lengths) of three PEXEL proteins as well as a construct containing the first 62 residues of the known PV-resident protein SERA5 whose signal peptide cleavage site was previously determined (35). All were tagged C-terminally with eGFP (Fig. 1A, S1). We integrated them for expression at the *attB* locus in the *P. falciparum* NF54*^attb^* strain (36). To determine their localization, we harvested 30h old parasite-infected RBCs and carried out a cell fractionation/anti-GFP western blot strategy (Fig. 1B).

**Figure 1:**
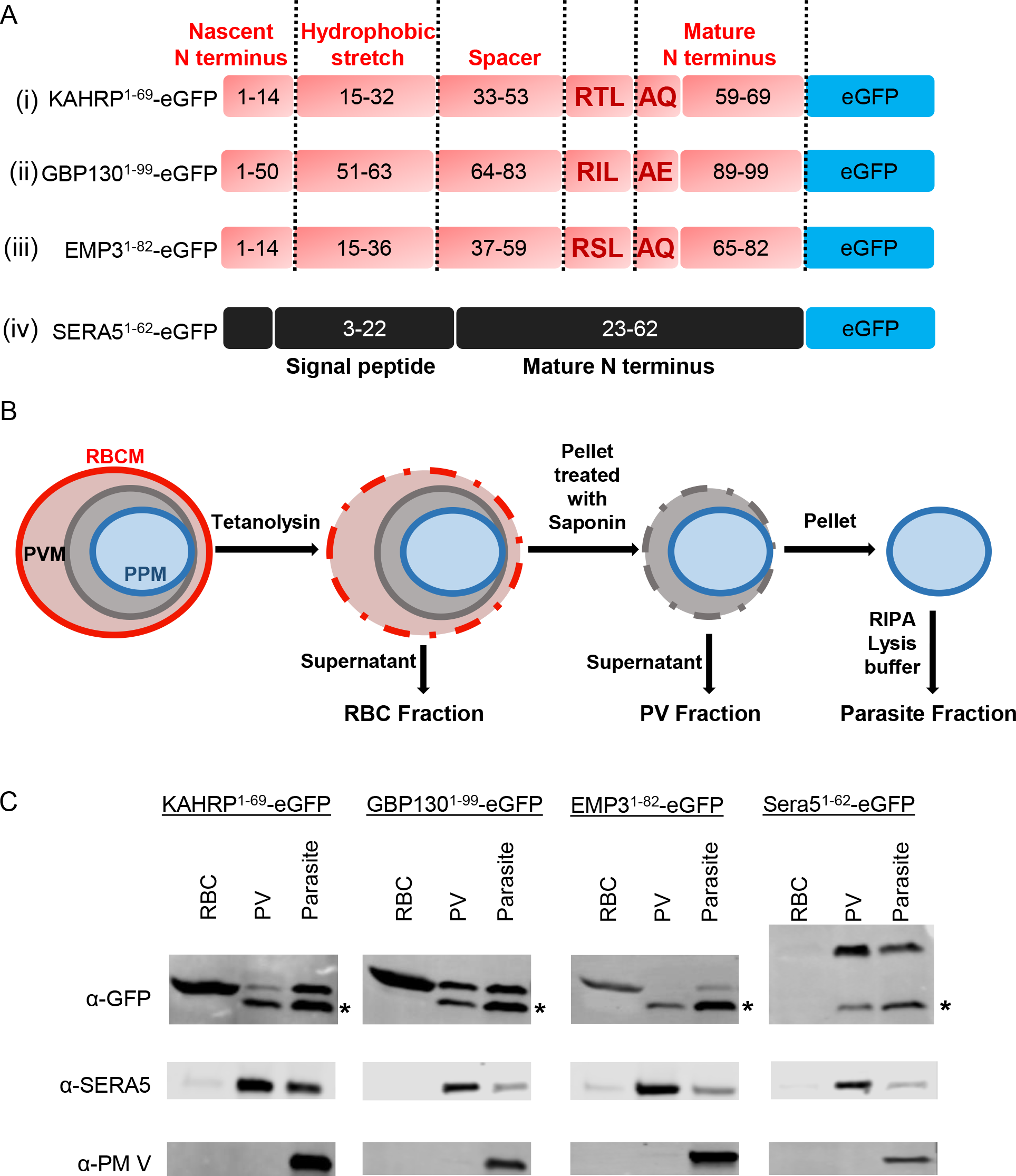
Minimal reporter constructs and their localization. (A) Schematic representation of minimal reporter constructs. (i) (ii) and (iii) are PEXEL reporters and (iv) is a PV resident reporter. Different segments of PEXEL reporters are delineated by dashed lines and labelled at the bottom for (iv). Residue numbers starting from the nascent N terminus are printed within each segment and the PEXEL motif residues are highlighted in bold red. (B) Compartment fractionation strategy. RBCM: Red blood cell membrane; PVM: Parasitophorous vacuole membrane; PPM: Parasite plasma membrane. (C) Representative western blots of different fractions from RBCs infected with *P. falciparum* expressing the reporters from A. Primary antibodies that were used to probe the blots are labelled at the left. The bottom band (marked with asterisks) in the anti-GFP blots is the free GFP band devoid of mature reporter portions. Each construct was tested at least 3 times.

Cells were treated with tetanolysin, a bacterial toxin that selectively perforates the RBC membrane (32, 37). Following centrifugation, we collected the supernatant as the infected RBC cytoplasm fraction and further treated the pellet with saponin. Saponin permeabilizes the PV membrane more efficiently than the PPM (38, 39). Supernatant from this treatment was collected as the PV fraction and the pellet was then fully lysed in RIPA lysis buffer as the intracellular parasite fraction. Western blotting of these fractions revealed the efficient export of the mature PEXEL reporters into the RBC and the retention of the SERA5 reporter in the PV (Fig 1C). We also probed the fractions with an anti-SERA5 antibody that does not recognize the reporter fragment (40). It revealed an enrichment of the native protein in the PV. The ER protein PM V was only detected in the intracellular parasite fraction (Fig 1C). These two proteins were also probed in subsequent experiments as markers of consistent fractionation. In our blots, we saw substantial free GFP in the intracellular parasite and PV fractions (marked with asterisks in Fig. 1C). This steered us away from using fluorescent microscopy and cautioned against the quantitative overinterpretation of previous studies.

### Mature N-terminal sequences of PEXEL proteins are sufficient for their export into the RBC

To test the necessity of the PEXEL motif for export, we constructed fusion reporters of SERA5 and KAHRP, in which modules N-terminal to the cleavage site are combined with modules C-terminal to the cleavage site (Fig. 2A, S1). The mature KAHRP reporter was efficiently cleaved (presumably by SP) and exported when placed after the SERA5 signal peptide (Fig. 2B). This suggests that its export signal resides C-terminal to the cleavage site and that neither the PEXEL motif nor processing specifically by PM V is required for export. On the other hand, the mature SERA5 reporter placed after the KAHRP RxL was cleaved (presumably by PM V), yet the conserved PEXEL residues were not sufficient to turn a PV resident reporter into an exported reporter. Therefore, a properly placed RxL is sufficient for cleavage in the ER but not for export. Taken together, these results demonstrate that if a protein travels to the PV from the ER, its mature N terminus determines its export capacity and not the pre-cleavage sequence.

**Figure 2:**
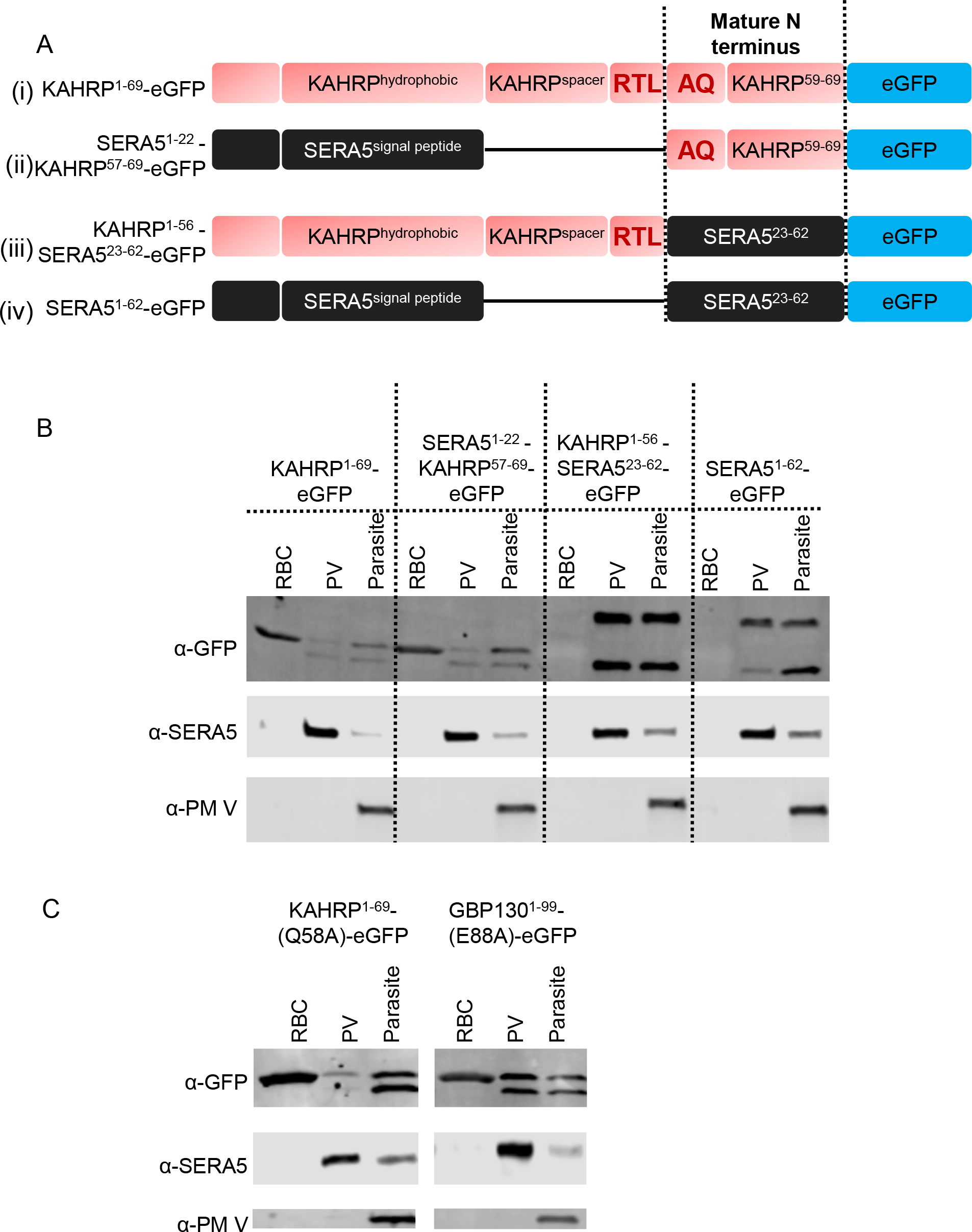
Investigation of the role of the PEXEL motif in protein export to the RBC. (A) Schematic representation of KAHRP and SERA5 fusion reporters (ii and iii). Original reporters (i and iv) are also shown for reference. (B) Representative western blots of different fractions from RBCs infected with *P. falciparum* expressing the reporters from A. Primary antibodies that were used to probe the blots are labelled at the left. The experiment was performed twice. (C) Representative western blots of different fractions from RBCs infected with *P. falciparum* expressing KAHRP and GBP130 reporters with alanine substituted semi-conserved 5^th^ positions of the PEXEL motif. Each construct was tested twice.

It can be argued from the above experiments that mature PEXEL reporters were exported due to the remnant part of the PEXEL, i.e. the xE/Q/D sequence at their N termini and the mature SERA5 reporter (starting with TG) was not exported because it did not have this signature at its N terminus (Fig. S1). We tested the importance of the semi-conserved 5^th^ residue of PEXEL by substituting it with alanine in both KAHRP and GBP-130 reporters. These substitutions did not inhibit their export, indicating that the 5^th^ residue of PEXEL is not required for RBC export (Fig. 2C).

### Substituting the hydrophobic stretch of the EMP3 reporter with the SERA5 signal peptide restores export

Previously, a case was made in favour of the essentiality of the PEXEL motif for export with the observation that the export of the minimal mature EMP3 reporter was abrogated when residues between the hydrophobic stretch and the mature N terminus of EMP3, including the RxL, were replaced with a single alanine. From this construct, the same mature reporter was produced, presumably by the action of SP after the alanine at the end of the hydrophobic stretch, yet the reporter was not exported (26). As this result is paradoxical to our conclusion, we decided to test a similar EMP3 reporter ourselves (Fig. 3A(ii), S1) and found, as previously observed, that the reporter did not pass the PV (Fig. 3B). Interestingly though, the level of the total mature reporter in this deletion construct was significantly lower compared to the original reporter (Fig. 3C). We then fused the mature EMP3 reporter after the SERA5 signal peptide (Fig. 3A(iii)). The total quantity and the export of the mature reporter from this construct were comparable to those of the full EMP3 reporter (Fig. 3C). Using mass spectrometry, we confirmed that the N termini of all three of these mature reporters were identical (Fig. S2).

**Figure 3:**
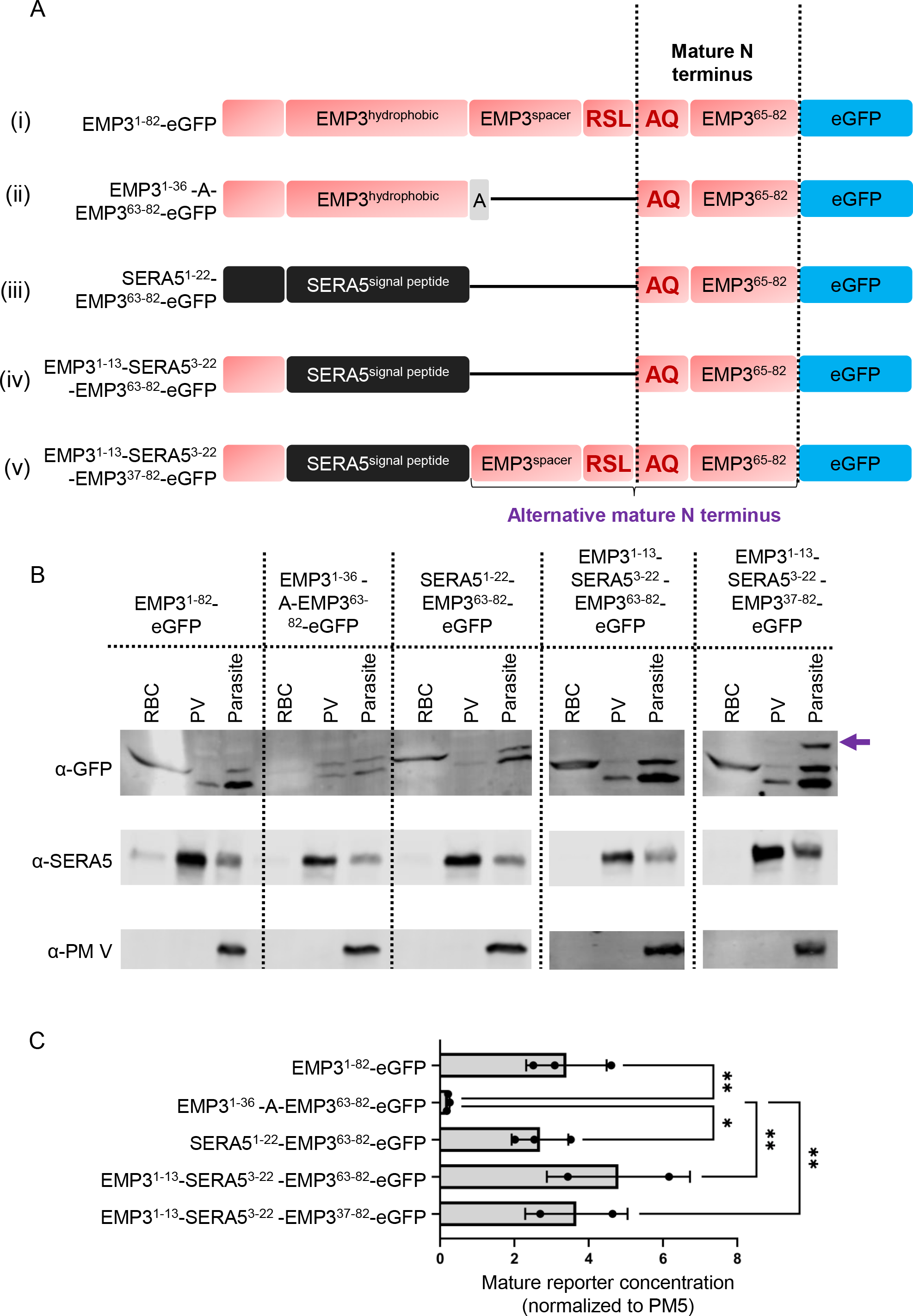
Processing, export and comparison of the protein level of different EMP3 reporters. (A) Schematic representation of the EMP3 reporter constructs. An alternative mature N terminus for the last construct is marked at the bottom (B) Representative western blots of different fractions from RBCs infected with *P. falciparum* expressing the reporters from A. Primary antibodies that were used to probe the blots are labelled at the left. Note the presence of an alternative mature form (marked with a purple arrow) for the last reporter in the parasite fraction. (C) Normalized western blot quantification of the standard mature forms of the reporters. Signals were combined from each fraction and then normalized to the PM V signal from the parasite fraction. * denotes a P ≤ 0.05 and ** denotes P ≤ 0.01 in Fisher’s LSD test. The P-value for the one-way ANOVA was 0.0113. Mean and standard deviations from 2 or 3 biological replicates are shown along with individual data points.

Efficient export of the SERA5^1-22^-EMP3^63-82^-eGFP fusion reporter again confirmed that the PEXEL motif is not essential for export and we hypothesized that the perplexing lack of export of the EMP3^1-36^ -A-EMP3^63-82^-eGFP construct had arisen from an inefficient cleavage after the hydrophobic stretch by the SP, which led to the degradation of the reporter. Protein level and export of the mature reporter were maintained when the nascent N-terminal residues of SERA5 were replaced with those from EMP3 (Fig. 3A (iv), B, C). When we did the same replacement in the full-length, PEXEL-containing EMP3 reporter (fig 3A(v)), in addition to the exported mature reporter, we also observed a higher molecular weight band in the parasite fraction whose size corresponds to an alternative mature reporter starting from the SP cleavage site (Fig. 3B, C).

Because we do not observe this higher molecular weight band in the original full-length reporter, it supports the argument that the signal cleavage site of EMP3 is non-functional. In any case, it was clear from our tested EMP3 reporters that the export deficiency of the mature reporter from the EMP3^1-36^ -A-EMP3^63-82^-eGFP construct could be reversed by changing the hydrophobic stretch of EMP3 without reintroducing the PEXEL motif.

### Putative structures of PEXEL protein mature N-termini resemble each other and are different than those of PV-resident proteins

As the RBC-export signal of the PEXEL proteins seemingly resides at their mature N terminus, we searched for commonalities in the first 10 amino acids following the cleavage sites of several PEXEL proteins for which there was experimental proof of RBC export (Fig. 4A, Table S1). As expected, there was barely any primary sequence conservation, except for the semi-conserved second position, which we already found to be not essential for export (Fig. 2C). We then looked at the AlphaFold structural predictions for that region of the selected proteins. Interestingly, most of them are predicted to form an alpha-helical structure (Fig. 4B, Table S1), albeit with low confidence scores. On the other hand, when we looked at the structures of the cognate residues following the signal peptide cleavage sites of a few known soluble PV resident proteins (Fig. 4C, Table S1), the majority of them had predicted random coil structures (Fig. 4D, Table S1). Based on this, we hypothesized that an alpha-helical structure at the mature N terminus of the PEXEL proteins is required for their export.

**Figure 4:**
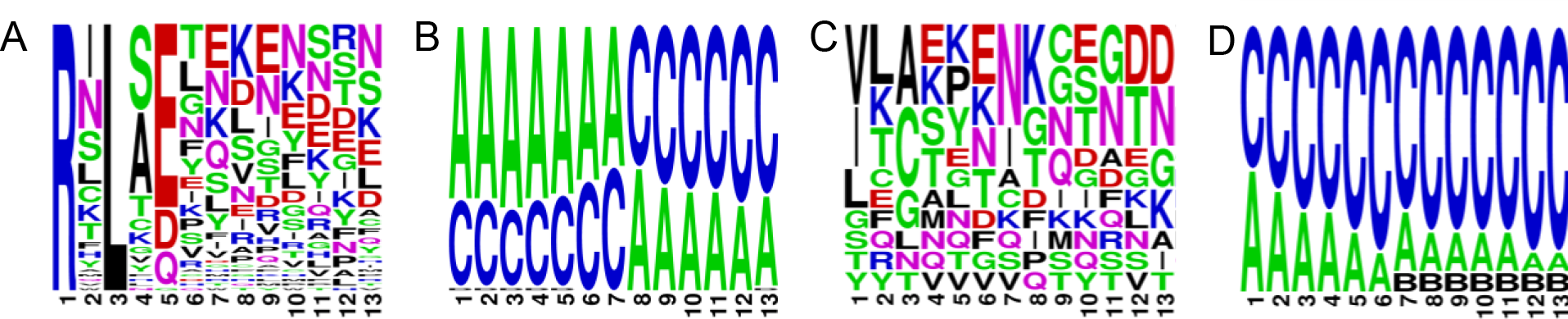
Primary sequences and putative structures of PEXEL and PV-resident proteins. (A) Amino acid frequency plots of 59 experimentally validated exported PEXEL proteins starting from the first position of the PEXEL motif to the 10^th^ position of the mature N terminus. (B) Frequency plot of AlphaFold structural predictions of the same residues shown in panel (A), where “A” denotes alpha-helical, “B” denotes beta-sheet and “C” denotes random coil. (C) Amino acid frequency plots of 13 experimentally validated PV-resident proteins starting from the -3 position of the signal peptide cleavage site to the 10^th^ position of the mature N terminus. (D) Frequency plot of AlphaFold structural predictions of the same residues shown in panel (A), with the letters denoting the same structural conformation as in panel (B).

### Helix-breaking proline insertion did not abrogate the export of reporters to the RBC

To test this hypothesis, we designed minimal reporter constructs of KAHRP and EMP3 where we inserted proline at the 3^rd^ or 6^th^ position of the mature reporters (Table S1). However, these insertions did not abrogate the export of the reporter constructs (Fig. 5A). Although proline is known to break alpha-helical structures, it might not work as such in the context of the KAHRP and EMP3 mature N terminal sequences (41–44). Therefore, we tested two other reporters, one where we placed a known alpha-helical sequence after the PEXEL cleavage site and another where a proline insertion has been experimentally determined to break the alpha helix conformation (Fig. S1) (45, 46). Using CD spectrometry, we verified the supposed conformations of these two peptides in vitro (Fig. 5B). However, in vivo, both of these reporters were exported into the RBC with similar efficiency (Fig. 5C, S1).

**Figure 5:**
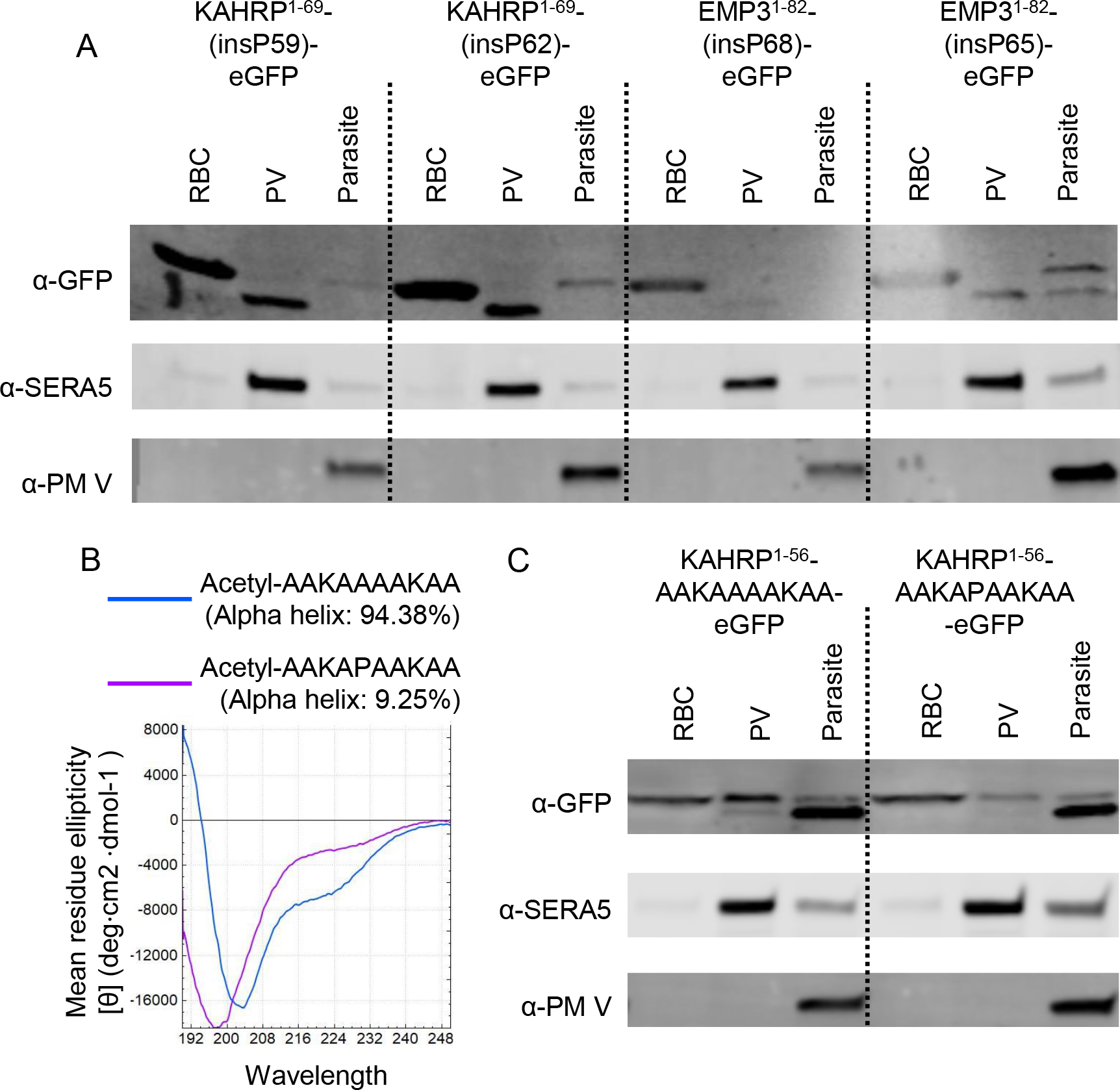
Investigation of the role of alpha-helical mature N terminus in protein export. (A) Representative western blots of different fractions of the KAHRP and EMP3 reporters with proline insertions at the 3^rd^ or the 6^th^ position of their mature N terminal region. The experiment was performed twice. (B) CD spectra of two small peptides, one of which takes an alpha-helical conformation in vitro whereas the other one loses the conformation due to alanine to proline substitution. (C)Representative western blots of different fractions from RBCs infected with *P. falciparum* expressing two artificially designed PEXEL reporters with the mature N terminal sequences shown in the construct name. The experiment was performed twice.

## Discussion

In this study, we have reevaluated the function of the PEXEL motif as the signal for the export of *Plasmodium* proteins into the RBC and found that it is not a direct prerequisite for export. We tested two PEXEL reporter constructs that conferred export as long as their export-competent mature N termini were exposed either by SP cleavage or by PEXEL cleavage. On the other hand, introducing a PEXEL cleavage site in a PV resident reporter allowed proper cleavage without altering its PV localization.

We also tested a previously published reporter (Fig. 3A(ii)) whose export was blocked upon removal of the PEXEL motif. We made variants of this reporter to test if the absence of the PEXEL motif was the cause of its export deficiency and found that replacing its ER targeting signal remedied the defect. The addition of the SERA5 signal peptide before the mature part of this reporter (Fig. 3A(iv)) possibly directed it to the SP-containing ER translocon, whereas the EMP3^1-13^-SERA5^3-22^ -EMP3^37-82^-eGFP ((Fig. 3A(v)) construct was likely targeted to both PM V and SP-containing translocons, maturing into two alternative reporters. In the absence of both SP and PM V cleavage sites, the EMP3^1-36^ -EMP3^63-82^-eGFP (Fig. 3A(ii)) construct was likely degraded by Endoplasmic Reticulum Associated Protein Degradation (ERAD) or proteasomal- degradation pathway. It is not completely clear why the small amount of the mature reporter liberated from this construct was not exported despite having an export-competent mature N terminus. We suspect that it might be related to the ER entry of the reporter because the knock- down of PfSPC25 and PfSec62, two components of the ER translocon required for protein import into the ER, also exhibited a decrease in protein levels for some native PEXEL proteins (4). Overall, more pertinent to the scope of this study, we showed that export could be re- established by introducing a functional SP cleavage site in this PEXEL-less reporter.

In light of these observations, two obvious questions emanate. One, if the PEXEL motif is not directly involved in protein export to RBC, what is its function and why is it conserved in so many exported proteins? And two, what is the real signal for export in the PEXEL proteins? The experiments presented here and other published experiments indicate that the PEXEL motif serves as a very potent cleavage signal at the ER of the parasite (26, 27, 30). We have shown that the hydrophobic stretch of the PEXEL protein EMP3 lacks a strong signal cleavage site, and that is also the case for several other PEXEL proteins tested before. For example, removing the RxL from PEXEL protein KAHRP results in more full-length reporter accumulation than for reporters cleaved after the hydrophobic stretch (5, 32). Another PEXEL protein HRPII accumulated as the full-length protein when PM V activity was inhibited (27). Pharmacological inhibition of PM V resulted in the accumulation of full-length EMP3 reporters, which strongly supports our conclusion (47, 48). Therefore, we think the PEXEL motif can be considered as a specialized signal cleavage site located distally from a hydrophobic signal anchor sequence of the PEXEL proteins and PM V acts as a non-canonical signal peptidase in *Plasmodium*. As the exported proteins are first loaded into the ER, it is unsurprising that a lot of them possess this motif.

In our attempt to address the second question, we searched for commonalities at the mature N terminus of several experimentally validated exported PEXEL proteins by analyzing their AlphaFold structure. We found an abundance of alpha-helical conformational predictions, which were absent from the corresponding sequences of several PV-resident proteins. A huge caveat of this analysis is that this structural prediction is based on the whole protein sequence where the mature N terminus is not yet liberated. Though the alpha helix is a very common secondary structure in proteins and not all the exported N termini had alpha-helical predictions, we decided to test its worth as the export signal because it is clear from the nonconserved nature of the primary sequences of exported N-termini that the export signal would not be a very unique one. Our results argue against the hypothesis that a simple alpha-helical structure at the mature N terminus would suffice as an export signal. However, this approach needs further refinements, and the experimental determination of the structure of the mature N termini of multiple PEXEL proteins could be enlightening. N termini of several PEXEL-negative exported proteins (PNEPs) also function as efficient export signals, indicating a common mechanism of selection of exported proteins (34). The role of N terminal acetylation is another interesting potential export signal. However, there are also examples of N terminally acetylated PV resident reporters (33), which indicates that it is not sufficient in itself as an export signal. Our data also supports this view as our export-deficient mature EMP3 reporter was acetylated. For now, the real signal for export is still unknown except that it is very promiscuous at the level of primary sequence and functions as an export signal only in the context of the N-terminal end of a protein.

As discussed in the introduction, multiple models have been proposed to connect two events in PEXEL protein trafficking, the cleavage by Plasmepsin V at the ER and the export through the PTEX channel at the PV membrane. Our conclusion indicates that these two events are independent of each other. Therefore, at least theoretically, there is no need for special sorting or chaperoning of the export-destined proteins from the ER; the selection can take place at the PV in its entirety. In the simplest scenario, an export-competent N terminus is recognized and differentiated from the export-incompetent N termini of PV-resident proteins by the PTEX complex (Fig. 6).

**Figure 6:**
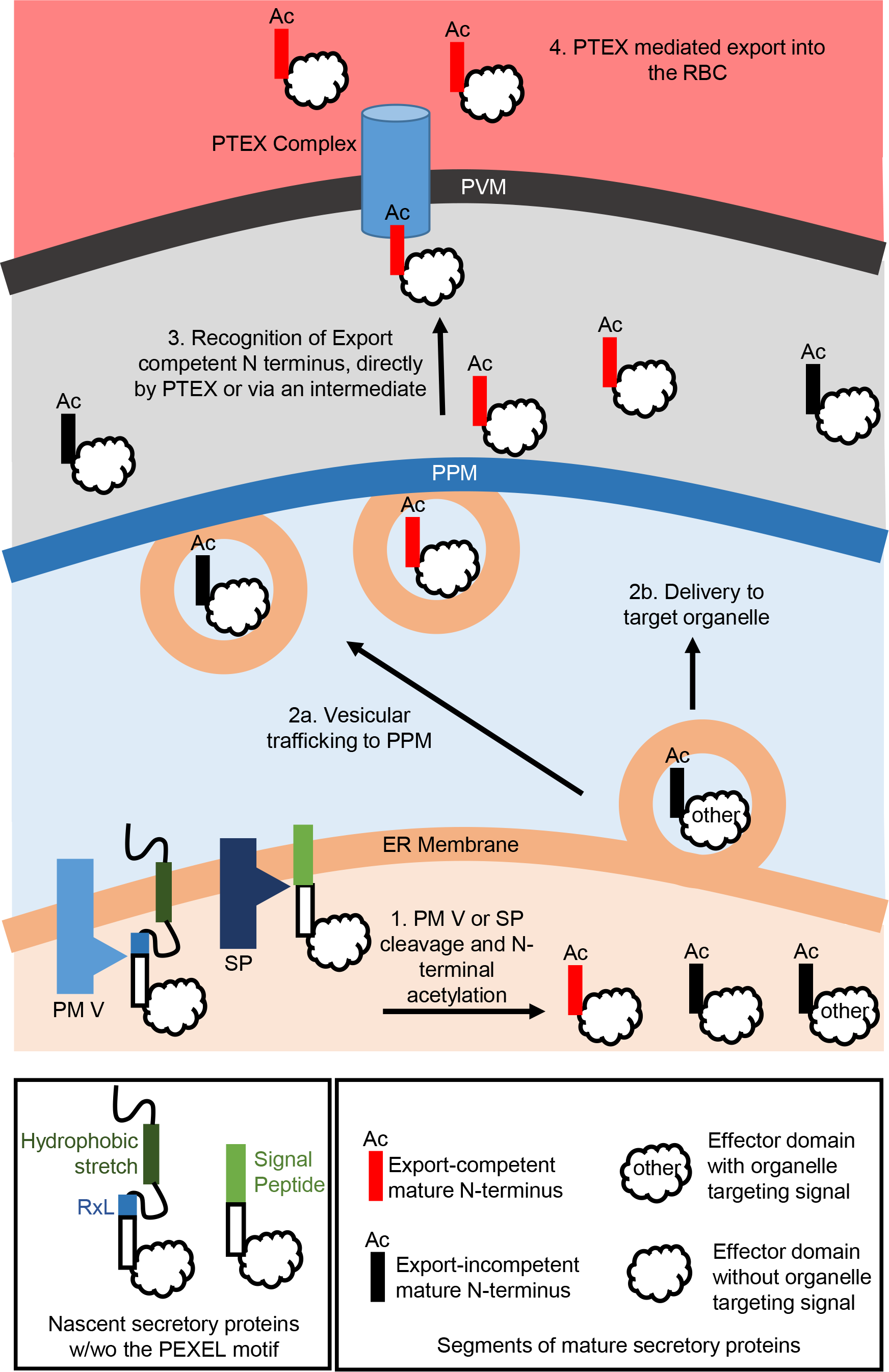
A model showing the trafficking of *P. falciparum* secretory proteins. Secretory proteins are targetted to the ER by their hydrophobic stretch and then cleaved by PM V at the PEXEL motif or by SP at the signal cleavage site, followed by acetylation. The cleavage liberates mature N termini that can be export-competent (shown in red) or incompetent (shown in black). There might be other organelle-targeting signals also in the mature proteins that direct them to their respective target organelle. Mature proteins secreted into the PV are recognized and loaded into the PTEX translocon based on the export competency of their mature N terminus.

Another important implication of our conclusion is that there might be PEXEL proteins that do not get exported into the RBC, but rather travel to other compartments from the ER. Examples of such proteins are rare but not nonexistent. For example, RESA is a dense granule protein that is cleaved by PM V in vitro and in vivo (23, 49, 50). Plasmepsin IX has an appropriately located PEXEL motif but localizes to the rhoptries (51). Therefore, it is important to experimentally determine the localization of PEXEL proteins rather than assuming that they are exported.

## Materials and Methods

### Maintenance of parasite cultures

*P. falciparum* strain NF54^attB^ was cultured in RPMI1640 (Gibco) media supplemented with 0.25% (w/v) Albumax (Gibco), 15 mg/l hypoxanthine, 110 mg/l sodium pyruvate, 1.19 g/l Hepes, 2.52 g/l sodium bicarbonate, 2 g/l glucose, and 10 mg/l gentamicin. Hematocrit concentration was maintained at 2%. Parasites expressing the reporters were maintained in 5nM WR99210. Human RBCs were collected from St Louis Children’s Hospital blood bank. Cultures were kept inside gas (5% O2, 5% CO2, and 90% N2) chambers at 37°C.

### Plasmid construction and transfection

A donor plasmid containing an attP site integrates into the *cg6 attB* locus of the NF54^attB^ strain when co-transfected with the pINT plasmid coding for Bxb1 integrase (52). All our reporter constructs were integrated into the genome using this strategy. They were expressed under the control of HSP86 (PF3D7_0708400) promoter and 3’UTR.

We first cloned the eGFP sequence between the AvrII and EagI sites of the pEOE-attP vector (53) using In-Fusion cloning (Clontech). KAHRP, GBP130, EMP3 and SERA5 minimal regions were amplified from NF54 mRNA isolated with TRIzol (ThermoFisher) using the SuperScript RT-PCR kit (Invitrogen). They were cloned into the XhoI-AvrII site of the pEOE-attP-eGFP vector using In-Fusion cloning. All the fusion constructs were made from the appropriate backbone vector with the QuikChange Lightning Multi Site Directed Mutagenesis kit (Agilent Technologies). Reporters were sequenced from Genewiz before transfecting the parasites. All the primers used for cloning and sequencing were purchased from IDT and their sequences are listed in Table S2.

Plasmids were isolated from bacterial clones using Nucleobond Xtra Midi (MN) kit and electroporated into the parasite as previously described (53). Successfully integrated clones were selected with media containing 5nM WR99210 from 36h post-transfection onwards as the donor plasmid codes for human dihydrofolate reductase (hDHFR) as the selection marker (54).

### Culture synchronization

An asynchronous parasite culture was washed in RPMI medium and then passed through a MACS LD magnet column (Miltenyi Biotec). NF54^attB^ parasites complete a replication cycle in 44 to 48h under our culture condition and older parasites (>28h old) are captured on the column due to the presence of paramagnetic hemozoin crystal (55). They were eluted in a prewarmed 2% hematocrit culture and incubated for 3 hours for egress and invasion. This culture was then treated with 5% sorbitol at 37°C for 10 minutes to osmotically lyse older parasites (due to the establishment of the new permeability pathway), leaving the newly invaded rings intact (56, 57).

Synchronized parasites were thereafter maintained by constantly shaking at 80 RPM under 5% parasitemia to maintain the synchrony.

### Compartment fractionation

Around 30h old synchronous parasite cultures were passed through the magnetic columns to harvest only infected RBCs. This step was critical because otherwise, haemoglobin from uninfected RBCs mask the western blot signals from a sample. Infected RBCs were washed twice in PBS and then treated with 50HU tetanolysin (Biological Laboratory Inc) in 60μl PBS plus HALT-Protease Inhibitor (PI) Cocktail (Thermo Fisher Scientific) for 10 minutes at room temperature. Following centrifugation at 1500g for 2 minutes, the supernatants were collected as the RBC fractions. The pellets were washed twice in PBS before treating with 60μl of 0.035% saponin in PBS-PI for 5 minutes on ice. Supernatants were collected as PV fractions and the pellets were washed twice before adding 60μl RIPA lysis buffer with PI. These were rapidly frozen and thawed using liquid nitrogen and a 42°C water bath three times and the supernatants were collected as the parasite fractions after 15 minutes of centrifugation at 4°C for 10 minutes. 20μl of 4X sample buffer with β-mercaptoethanol as the reducing agent was mixed with each sample and boiled for 5 minutes before storage at -20°C.

### SDS PAGE and western blotting

15 μl samples from each fraction were run in a 4-15% gradient gel (Biorad) and then transferred to a PVDF membrane for western blotting. We used mouse anti-GFP (Takara) at 1:1000 dilution, mouse anti-PM V (58) at 1:250 dilution and rabbit anti-SERA5 (40) at 1:1000 dilution as primary antibodies and IRDye conjugated goat secondary antibodies (LICOR) at 1:15000 dilution. Blots were incubated with primary antibodies O/N at 4°C and with secondary antibodies for 1 hour at room temperature. Licor Odessey blocking buffer was used for blocking and primary antibody dilutions and PBS plus 1% tween-20 was used to prepare secondary antibody dilutions as well as in all the washing steps. Blots were imaged in a Licor Odyssey imager and images were prepared (and quantified when required) using Image Studio Lite 5.2 (Licor). The protein level of EMP3 reporters was calculated by adding the intensities of reporter GFP bands (excluding the free GFP or any higher molecular weight band) from all three fractions and then normalizing that value for the PM V band intensity from the parasite fraction. Statistical analyses were performed in GraphPad Prism.

### Preparation of samples for mass spectrometry

Infected RBCs were harvested using magnetic columns from 300ml 5% parasitemia cultures and then directly lysed with 500ul GFP-trap lysis buffer (10 mM Tris/Cl pH 7.5, 150 mM NaCl, 0.5 mM EDTA, 0.5 % Nonidet™ P40 Substitute) by freeze-thaw. Supernatants were incubated with GFP-trap magnetic agarose (ChromoTek) at 4°C for 1 hour with continuous rotation. The beads were washed 3 times with wash buffer (10 mM Tris/Cl pH 7.5, 150 mM NaCl, 0.05 % Nonidet™ P40 Substitute, 0.5 mM EDTA) and then 50 μl 2x sample buffer with β-mercaptoethanol was added to the beads and boiled for 5 minutes for elution. All the eluted samples were run in a Biorad gradient gel. Specific bands were visualized with Coomassie blue staining and cut out of the gel for submission to the Mass Spectrometry Technology Access Center.

### Proteomics and data analysis

The protein gel bands were subjected to in-gel digestion. Each gel band was washed in 100 mM Ammonium Bicarbonate (AmBic)/Acetonitrile (ACN), reduced with 10 mM dithiothreitol, and cysteines were alkylated with 100mM iodoacetamide. Gel bands were washed in 100mM AmBic/ACN prior to adding 1 µg trypsin for overnight incubation at 37°C. The supernatant containing peptides was saved into a new tube. Gel was washed at room temperature for ten minutes with gentle shaking in 50% ACN/5% FA, and the supernatant was saved to peptide solution. The wash step was repeated each by 80% ACN/5% FA, and 100% ACN, and all supernatant was saved and then subject to the speedvac dry. After lyophilization, peptides were reconstituted with 0.1% FA in water. Peptides were injected onto a Neo trap cartridge coupled with an analytical column (75 µm ID x 50 cm PepMap^TM^ Neo C18, 2 µm). Samples were separated using a linear gradient of solvent A (0.1% formic acid in water) and solvent B (0.1% formic acid in ACN) using a Vanquish Neo UHPLC System coupled to an Orbitrap Eclipse Tribrid Mass Spectrometer with FAIMS Pro Duo interface (Thermo Fisher Scientific).

The resulting tandem MS data was queried for protein identification against the custom database, Plasmodium falciparum 3D7 database plus the 3 custom proteins (The cleaved form of EMP3(i), (ii), and (iii)), using Mascot v.2.8.0 (Matrix Science). The following modifications were set as search parameters: peptide mass tolerance at 20 ppm, trypsin enzyme, 3 allowed missed cleavage sites, carbamidomethylated cysteine (static modification), and oxidized methionine, deaminated asparagine/glutamine, and protein N-term acetylation (variable modification). The search results were validated with 1% FDR of protein threshold and 90% of peptide threshold using Scaffold v5.2.1 (Proteome Software). Data are available via ProteomeXchange with identifier PXD041451. Please use the following credential for review: Username: Password: m25z5Zvm.

### CD Spectrometry

Custom peptides with N-terminal acetylation were purchased from Biomatik. Peptides were diluted in 10 nM potassium phosphate buffer with 40% TFE with a final concentration of 0.2 mg/ml. CD spectra were recorded on a JASCO-J715 polarimeter (JASCO, Tokyo, Japan) over the wavelength range 190-250 nm in a 1-mm path length quartz cuvette using a step size of 0.1 nm. For each wavelength, three scans were performed. AVIV software was used for background subtraction. Mean residual ellipticity [θ] vs Wavelength plots were generated using CDtoolX (59) and online server K2D3 (60) was used to determine the alpha-helical composition of the peptides.

### Analysis of PEXEL and PV-resident protein sequences

Experimentally validated exported PEXEL proteins were selected from the list of the PEXEL proteins from Jonsdottir TK et al. (1). If multiple proteins had the same primary sequence at their mature N termini (mainly from exported protein families like RESA, RIFIN, STEVOR etc.), only one sequence was included in the list. PV-resident proteins were manually selected by reviewing several publications. The relevant part of their protein sequences and AlphaFold structures were taken from the Plasmodb database (61). A, B and C were used as codes if a particular residue was part of an alpha-helix, beta-sheet or random coil structure. Frequency plots of the amino acids or structures were created using the Weblogo tool (62).

## Acknowledgements

Mass Spectrometry analyses were performed by the Mass Spectrometry Technology Access Center at McDonnell Genome Institute (MTAC@MGI) at Washington University School of Medicine. We thank the group of Dr Michael Blackman from The Francis Crick Institute for the anti-SERA5 antibody and the group of Dr Sergej Djuranovic from the department of Cell Biology & Physiology at Washington University School of Medicine for the eGFP construct. This work was supported by NIH grant AI17106201.

---

**Supplementary Figure 1:**
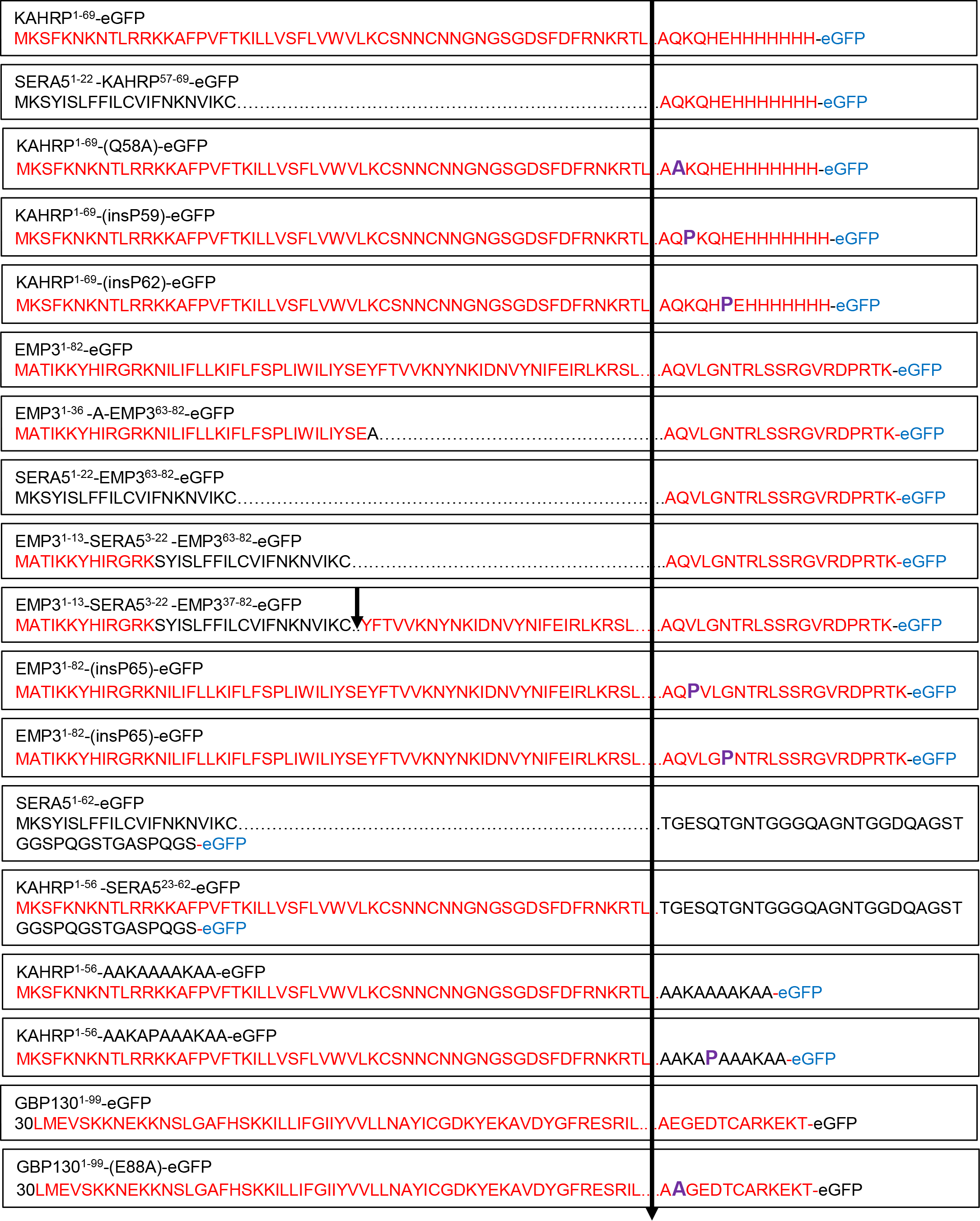
Full sequence of all the reporters in this study. eGFP sequence is not shown as well as the first 29 residues of GBP130. Bold arrows denote cleavage sites. Sequences from PEXEL proteins are printed in red. Substitutions and insertions are highlighted in bold purple.

**Supplementary Figure 2:**
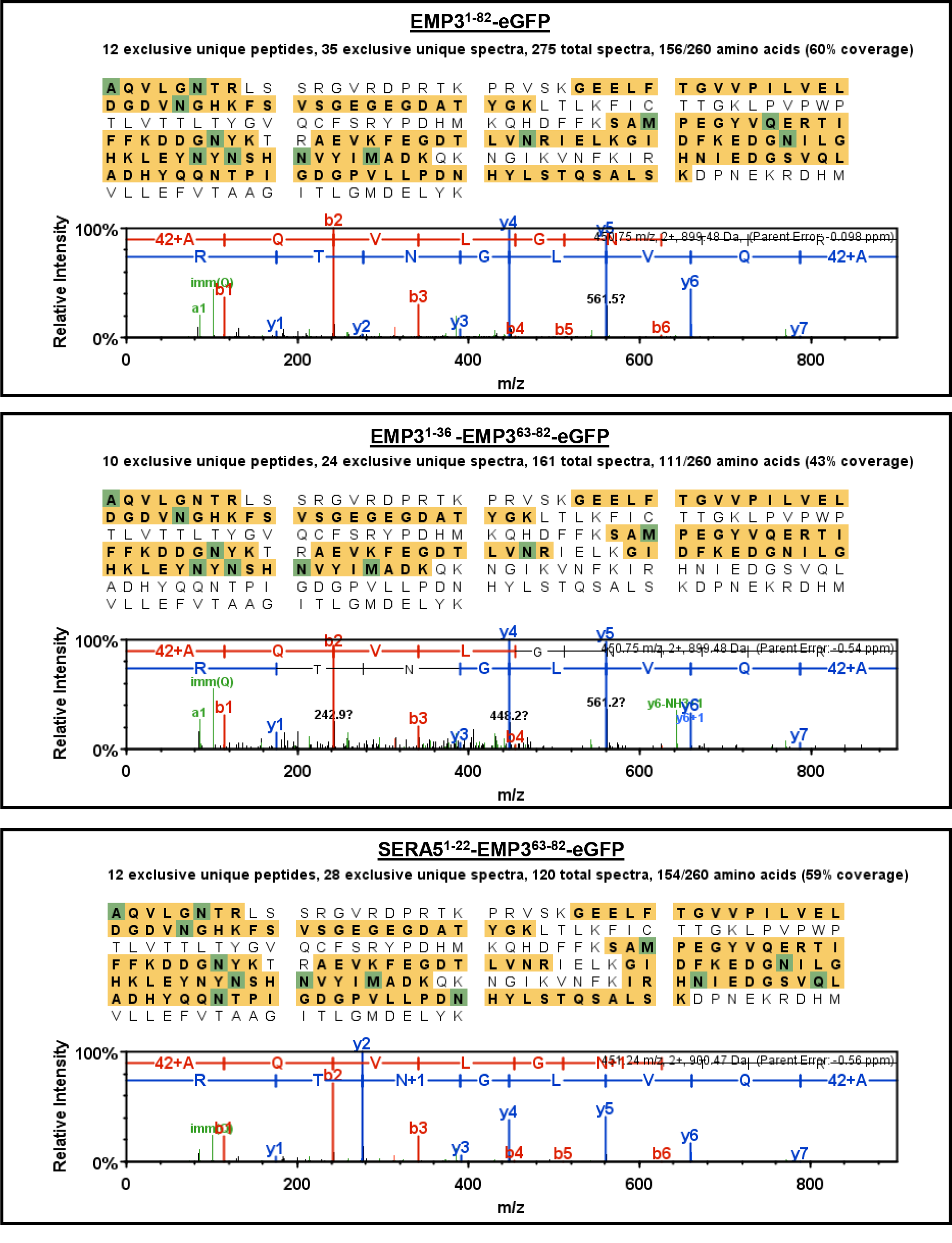
Coverage map and most N terminal peptide spectrum of EMP3 reporters. Construct names are on top. The coverage map highlights detected peptides at the 95% threshold. Green shades in the coverage map denote post-translational modification.

**Supplementary table 1: List of PEXEL proteins and PV resident proteins along with their mature N terminal sequences and AlphaFold structural predictions.** These sequences were used to generate the frequency plots shown in Figure 4. For the AlphaFold structural predictions, “a” denotes alpha-helical, “b” denotes beta-sheet and “c” denotes random coil.

Supplementary table 2: List of primers used in this study

